# UltraCortex: Submillimeter Ultra-High Field 9.4 T Brain MR Image Collection and Manual Cortical Segmentations

**DOI:** 10.1101/2024.06.18.599541

**Authors:** Lucas Mahler, Julius Steiglechner, Benjamin Bender, Tobias Lindig, Dana Ramadan, Jonas Bause, Florian Birk, Rahel Heule, Edyta Charyasz, Michael Erb, Vinod Jangir Kumar, Gisela E Hagberg, Pascal Martin, Gabriele Lohmann, Klaus Scheffler

**Affiliations:** Department High-field Magnetic Resonance, Max Planck Institute for Biological Cybernetics, Tübingen, Germany; Department of Biomedical Magnetic Resonance, University Hospital Tübingen, Tübingen, Germany; Department of Diagnostic and Interventional Neuroradiology, University Hospital Tübingen, Tübingen, Germany; Center for MR Research, University Children’s Hospital, Zurich, Switzerland; Department of Neurology and Epileptology, Hertie Institute for Clinical Brain Research, University of Tübingen, Tübingen, Germany

## Abstract

The *UltraCortex* repository (https://www.ultracortex.org) houses magnetic resonance imaging data of the human brain obtained at an ultra-high field strength of 9.4 T. It contains 86 structural MR images with spatial resolutions ranging from 0.6 to 0.8 mm. Additionally, the repository includes segmentations of 12 brains into gray and white matter compartments. These segmentations have been independently validated by two expert neuroradiologists, thus establishing them as a reliable gold standard. This resource provides researchers with access to high-quality brain imaging data and validated segmentations, facilitating neuroimaging studies and advancing our understanding of brain structure and function. Existing repositories do not accommodate field strengths beyond 7 T, nor do they offer validated segmentations, underscoring the significance of this new resource.

## Background & Summary

Here we introduce the *UltraCortex* repository (https://www.ultracortex.org), housing magnetic resonance imaging (MRI) data of the human brain obtained at an ultra-high magnetic field strength of 9.4 T. The repository comprises 86 *T*_1_-weighted structural MR images of healthy human brains, acquired at spatial resolutions ranging from 0.6 to 0.8 mm using MP-RAGE and MP2RAGE sequences. Additionally, UltraCortex includes segmentations of 12 brains into gray matter (GM) and white matter (WM) compartments. These segmentations have undergone independent validation by two expert neuroradiologists, thus establishing them as a reliable gold standard.

While repositories containing MRI data at 7 T exist^1–3^, none are currently available for field strengths beyond 7 T. Moreover, validated segmentations of UHF MRI data are presently lacking, but are urgently needed to support the development of new algorithms and tools for UHF MRI data analysis, and enable the validation of existing methods. It should be noted that we refer to field strengths *≥* 7 T as *ultra-high field* (UHF).

MRI has made a tremendous contribution to our understanding of the human brain. However, ultra-high resolution brain imaging is constrained by technical challenges. Image resolution and signal-to-noise ratio (SNR) both increase with field strength, resulting in improved image quality at a UHF of 9.4 T^4^. This makes it easier to visualize structures with low inherent signal, such as small lesions or areas of low proton density. The enhanced spatial resolution of UHF MRI is beneficial for capturing fine anatomical details where conventional resolutions of 1 – 2 mm are insufficient. The improved tissue contrast of UHF MRI allows for clearer classification of different tissue types, which is critical when studying various pathological conditions. In addition, UHF MRI offers enhanced sensitivity for functional MRI (fMRI), allowing more precise localization of brain activity and more detailed spectroscopic imaging, which is useful for studying metabolites in conditions such as Alzheimer’s disease and oncology.

The availability of large neuroimaging databases on open platforms such as OpenNeuro (https://openneuro.org), Image and Data Archive (https://ida.loni.usc.edu), and Human Connectome Project^5^ (https://humanconnectome.org) has greatly advanced the field of neuroimaging. These repositories have not only facilitated the development of imaging techniques, but have also provided a fertile ground for the development and application of new algorithms for the analysis of brain images.

However, the majority of existing data analysis algorithms were designed and optimized for MRI data acquired at lower field strengths, typically 3.0 T or 1.5 T. In structural imaging, a spatial resolution of 1 mm^3^ has become the standard. Consequently, when applied to UHF MRI data, many algorithms use downsampling, resulting in a loss of the beneficial enhancements provided by UHF MRI. This mismatch highlights a bottleneck in UHF MRI research: the underutilization of the detailed data that UHF MRI can provide, constrained by the limitations of existing analysis tools. The development and validation of new algorithms rely heavily on access to high-quality public databases, but there is currently no large public UHF MRI database with field strengths of greater than 7 T and resolutions of smaller than 1 mm. This becomes even more apparent when considering the availability of manually segmented MR images, the *gold standard* for anatomical segmentation. Only a small number of manually segmented images are available at ≤7T^3,6,7^, and none are available at 9.4 T. This limitation has significant implications for brain analysis, leading to inaccuracies in structural measures such as cortical volume, thickness, and depth, and thus potentially misleading results. Furthermore, the application of these imaging techniques to fMRI analysis is also affected, as errors in the initial segmentation phase can cascade throughout the analysis pipeline.

With UltraCortex we aim to provide a foundation for the development and validation of new algorithms for UHF MRI data analysis. It can be used to develop new algorithms that are applicable across different field strengths and resolutions, as well as to validate and benchmark existing algorithms. The data can also be used for image denoising, artifact removal, image quality enhancement and super-resolution. Additionally, the data can be utilized for educational purposes, such as to illustrate the intricate details of brain structures. The following sections describe the initial release of UltraCortex, including the data acquisition, processing and validation steps. Figure 1 summarizes the dataset.

**Figure 1.**
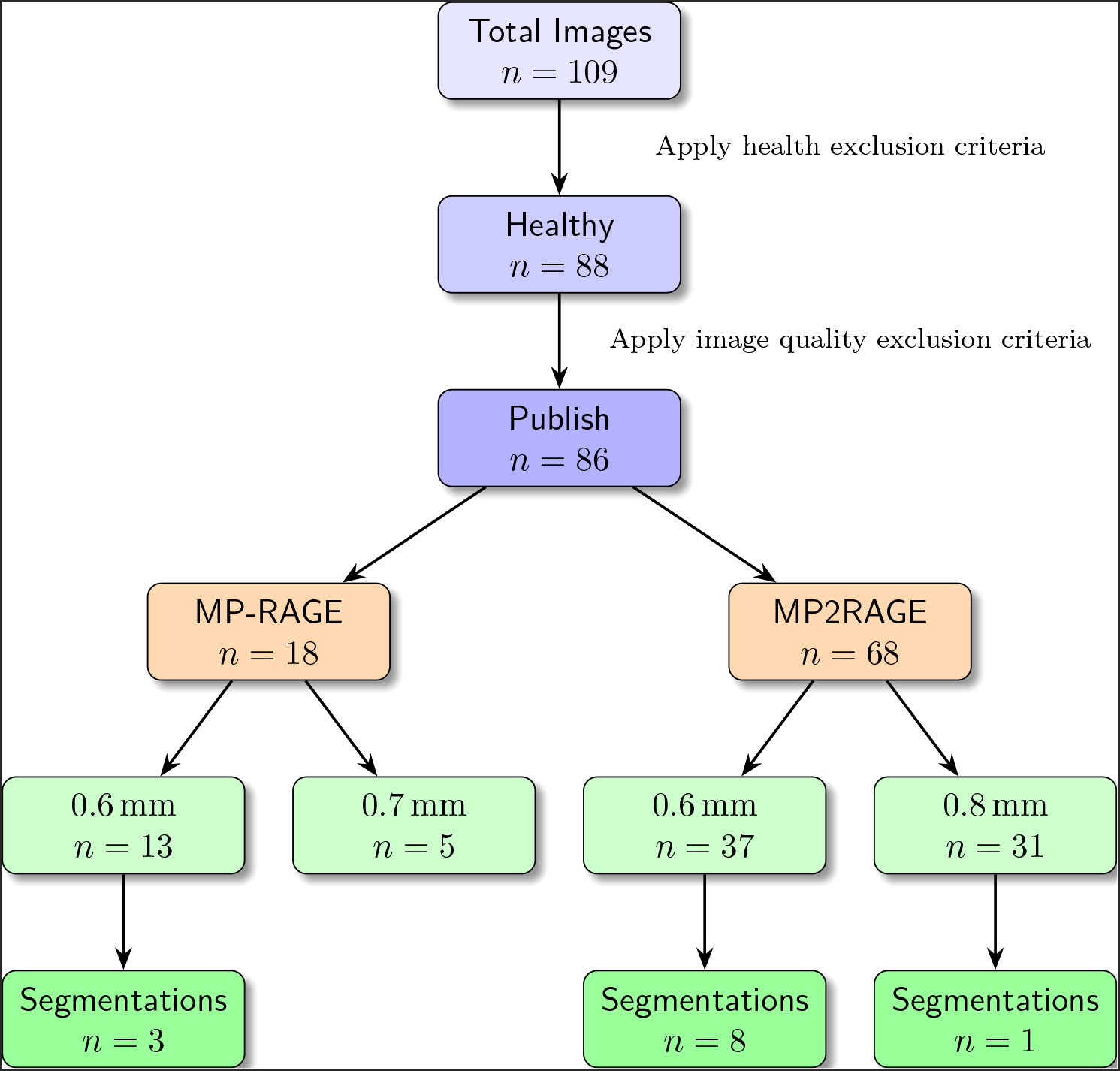
Dataset overview. The chart shows (i) from top to bottom how images were included or excluded for the subsequent steps of the pipeline, and (ii) the final distribution of the dataset across sequences, resolutions, and existing segmentations.

## Methods

### Demographics

Subsequent to the original measurements, we collected 86 *T*_1_-weighted structural brain MR images from 78 healthy adult volunteers (28 females and 50 males; age range: 20.0 – 53.0 years; median age at acquisition: 29.0 years; interquartile range of age: 7.8 years). 6 subjects were measured in two independent sessions and 1 subject was measured in three sessions. The demographic data, including participant sex and age at the time of scanning, are presented in Table 1. Some of the subject demographics and image data in this repository have been used in previous studies^8^.

**Table 1.**
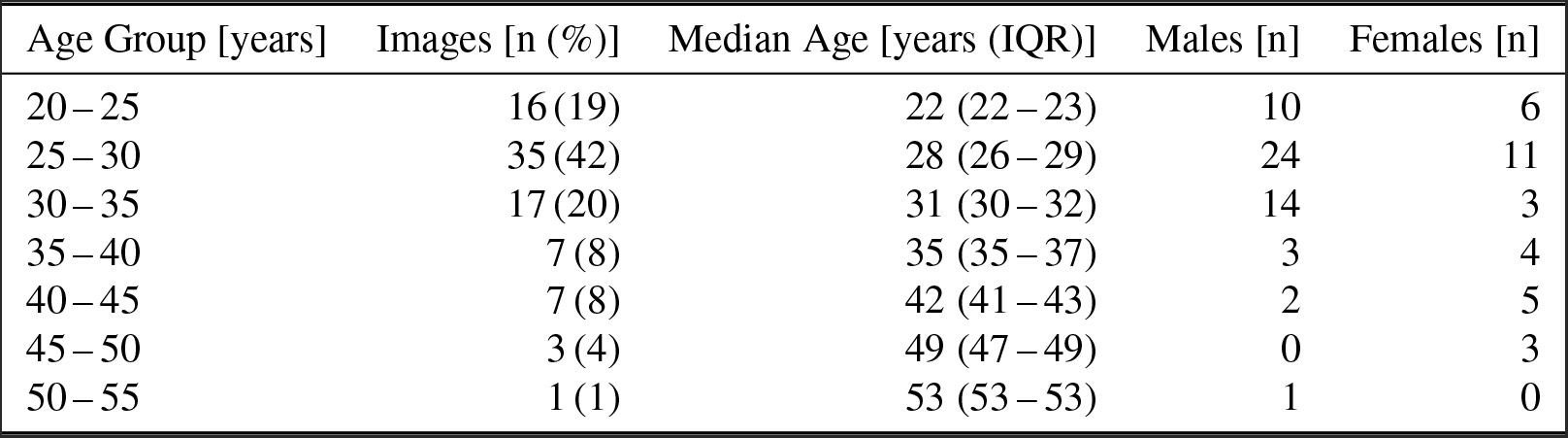
Age and sex characteristics of dataset.

Participants were selected in accordance with the guidelines and ethics of the Max Planck Institute for Biological Cybernetics, Tübingen, Germany and the University Hospital, Tübingen, Germany. Prior to enrollment in an MRI study, all subjects underwent a review of inclusion and exclusion criteria by a qualified physician. This included ensuring that, unless otherwise specified, there was no known history of neurological or psychiatric disease and that there were no contraindications to MRI scans. Additionally, all participants were required to have normal or corrected-to-normal vision. Prior to participation, written informed consent was obtained from all participants. This consent included an agreement for open sharing of anonymized data.

Due to the more rigorous clinical criteria employed in this dataset, 21 participants were excluded from an original collection of 109 images from 101 subjects. Furhter, 2 images were excluded due to a failed technical validation, resulting in the final dataset.

### Data Acquisition

Data acquisition was conducted using a 9.4 T whole-body MRI scanner (Siemens Healthineers, Erlangen, Germany) at the Department for High-field Magnetic Resonance of the Max Planck Institute for Biological Cybernetics. The scanner was equipped with a 16-channel dual-row transmit array, operating in CP+ (circularly polarized) mode, and paired with a 31-channel receive array^9^. After *B*_0_ shimming, whole-brain *T*_1_-weighted magnetization-prepared rapid acquisition gradient echo (MP-RAGE)^10^ or modified MP-RAGE (MP2RAGE)^11,12^images were acquired. Both sequences used a TR-FOCI pulse optimized for 9.4 T for magnetization inversion^12,13^. Nevertheless, despite the brief echo time used for the scans, signal dropouts due to strong *B*_1_-field non-uniformity close to the ear canals were observed in all subjects. It was determined that the data regarding the gradient nonlinearity profile would not be collected, indicating that the correction of gradient nonlinearity effects would not be conducted, as they play a relatively minor role within the field of view. It is important to note, however, that quantitative analysis, such as volumetric measurements, may be affected by this. The sequence parameters were empirically optimized for the contrast between GM and WM tissues. An example of typical images is illustrated in Figure 2. The parameter setups differ in sequence, number of acquired sagittal slices isotropic voxel size, acquisition matrix, inversion times TI1 and TI2, repetition time TR, echo time TE, flip angle FA, acceleration factor R of GRAPPA^14^ technology for parallel acquisition and reconstruction, slice partial Fourier factor PFF = 6*/*8, and optional CAIPI^15^ shift in the GRAPPA technology. An overview of the sequence configurations in use is presented in Table 2. The comprehensive image-specific scanning parameters can be accessed via the data records.

**Table 2.** Details of the dataset, outlining the various scanning parameter sets. MR Images acquired at the Department for High-field Magnetic Resonance of the Max Planck Institute for Biological Cybernetics, Tübingen, Germany.

| Sequence | Images | Voxel Size [mm<sup>3</sup>] | slices | matrix | TR [ms] | TE [ms] | FA [°] | R | CAIPI |
| --- | --- | --- | --- | --- | --- | --- | --- | --- | --- |
| MP-RAGE | 18 |  |  |  |  |  |  |  |  |
| | 4 | $0.7 \times 0.7 \times 0.7$ | 240 | $300 \times 306$ | 6500 | 3.4 | 7.5 | $2 \times 1$ | off |
| | 1 | $0.7 \times 0.7 \times 0.7$ | 256 | $300 \times 306$ | 6500 | 3.4 | 7.5 | $2 \times 1$ | off |
| | 1 | $0.6 \times 0.6 \times 0.6$ | 288 | $352 \times 352$ | 6000 | 2.3 | 8.0 | $2 \times 2$ | on |
| | 4 | $0.6 \times 0.6 \times 0.6$ | 288 | $352 \times 352$ | 3800 | 2.5 | 6.0 | $2 \times 2$ | on |
| | 8 | $0.6 \times 0.6 \times 0.6$ | 288 | $320 \times 320$ | 3800 | 2.5 | 6.0 | $2 \times 2$ | on |
| MP2RAGE | 68 |  |  |  |  |  |  |  |  |
| | 11 | $0.8 \times 0.8 \times 0.8$ | 192 | $256 \times 256$ | 9000 | 2.3 | 4.0 | $3 \times 1$ | off |
| | 20 | $0.8 \times 0.8 \times 0.8$ | 236 | $256 \times 256$ | 6000 | 2.3 | 4.0 | $3 \times 1$ | off |
| | 1 | $0.6 \times 0.6 \times 0.6$ | 256 | $352 \times 352$ | 6000 | 3.0 | 4.0 | $2 \times 2$ | on |
| | 6 | $0.6 \times 0.6 \times 0.6$ | 256 | $352 \times 352$ | 6000 | 3.0 | 5.0 | $2 \times 2$ | off |
| | 27 | $0.6 \times 0.6 \times 0.6$ | 256 | $352 \times 352$ | 6000 | 3.0 | 5.0 | $2 \times 2$ | on |
| | 3 | $0.6 \times 0.6 \times 0.6$ | 260 | $261 \times 315$ | 6000 | 3.0 | 5.0 | $2 \times 2$ | on |

**Figure 2.**
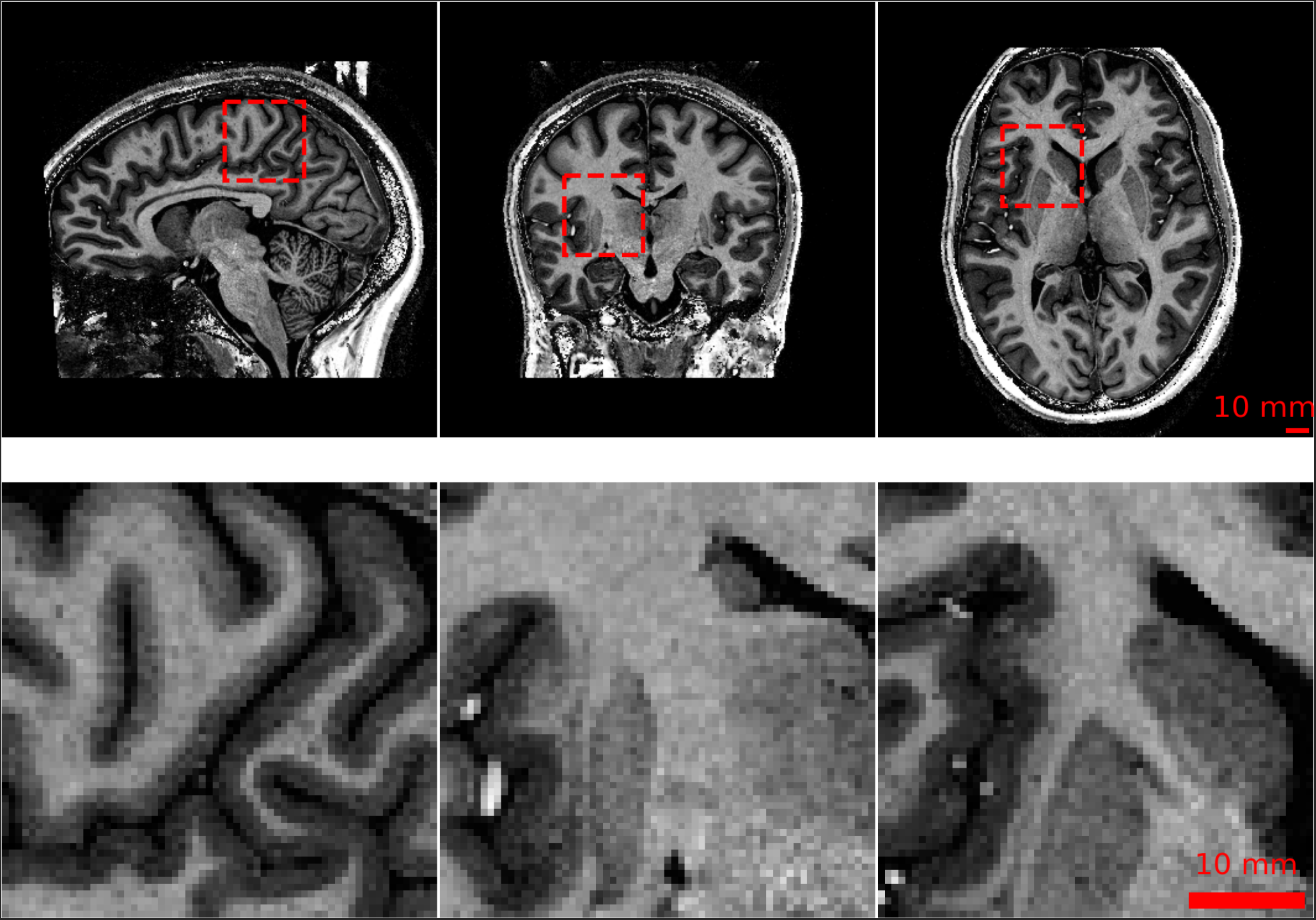
Example 9.4 T MRI slices from subject `sub-73` acquired with 0.6*×*0.6*×*0.6 mm^3^. The top row shows from left to right sagittal, coronal, and axial images and ROIs of the corresponding full resolution data in the bottom row.

### MP-RAGE

A total of 18 whole-brain MP-RAGE images were acquired with five different sequence parameter setups and two different resolutions: (i) 5 images with voxel size = 0.7 *×* 0.7 *×* 0.7 mm^3^ and (ii) 13 images with voxel size = 0.6 *×* 0.6 *×* 0.6 mm^3^.

### MP2RAGE

A total of 68 whole-brain MP2RAGE images were acquired with six distinct sequence configurations and two different resolutions: (i) 31 images with voxel size = 0.8 *×* 0.8 *×* 0.8 mm^3^ and (ii) 37 images with voxel size = 0.6 *×* 0.6 *×* 0.6 mm^3^.

### Processing

From each MP2RAGE scan, four distinct volumes were generated: INV1, INV2, UNI, and T1. The UNI volume represents a composite of the images acquired at both inversion times (TI1 and TI2), whereas the T1 volume is a *T*_1_ estimation map derived from the volumes at TI1 and TI2. Background noise was removed from the T1 volume using a regularization term during contrast division when the images from both inversion times were combined^16^. MP2RAGE images were reconstructed offline using customized routines that use the 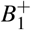 map when available to correct for the flip angle variation caused by the shorter wavelength at 9.4 T^12^. The remaining *T*_1_-weighted DICOM (https://www.dicomstandard.org) files were converted to NIfTI-1 (https://nifti.nimh.nih.gov) format using dcm2niix^17^. The NIfTI files were then anonymized by first using SynthStrip^18^ to extract the brain mask and then using Quickshear^19^ to de-identify the images. In addition, all images were stripped of any potentially revealing metadata in the NIfTI headers.

### Segmentation

We selected 12 subjects whose ultra-high field 9.4 T MRI scans were of high quality for detailed manual segmentation. The objective of the segmentation was to obtain high-precision masks of the cortical ribbon. In accordance with FreeSurfer^20^, the labels shown in Table 3 were segmented. This implies that subcortical structures, such as the basal ganglia, were added to the “inner” label, namely the cerebral white matter. However, the hippocampi and amygdalae were added to the background according to the FreeSurfer ribbon file. For cortical areas that exhibited signal dropouts, we opted to segment a topological closed cortex, leading to guessed labels in this region.

**Table 3.**
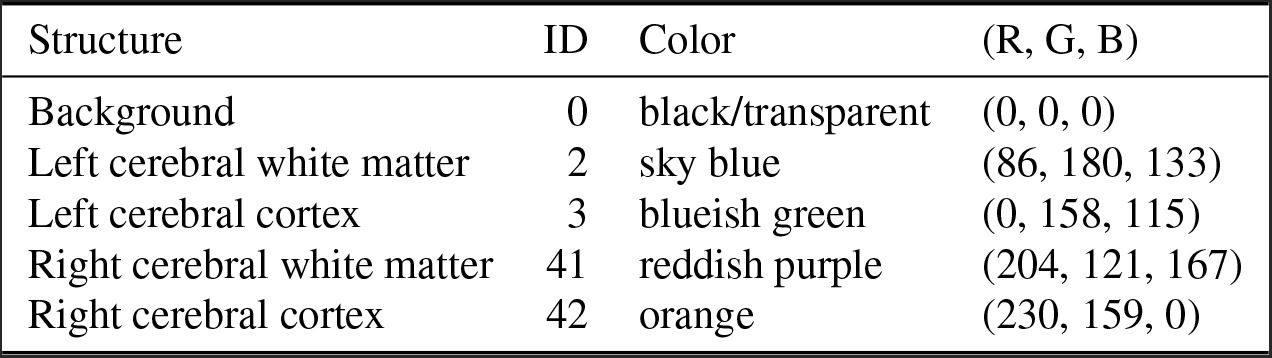
UltraCortex segmentation IDs and mapping to color scheme in Figure 3.

At a high level, the segmentation procedure can be described as follows: (i) FreeSurfer generation of surrogate segmentations, (ii) iterative manual corrections interleaved with expert neuroradiologist review, and (iii) final validation of the individual segmentations by two neuroradiologists. Given the distinctive characteristics and appearance of 9.4 T images, we sought to engage the expertise of neuroradiologists with extensive experience in interpreting UHF data acquired at 9.4 T. Figure 3 illustrates an examplar of the segmentation, while Figure 4 shows the average necessity for modifying voxel labels.

**Figure 3.**
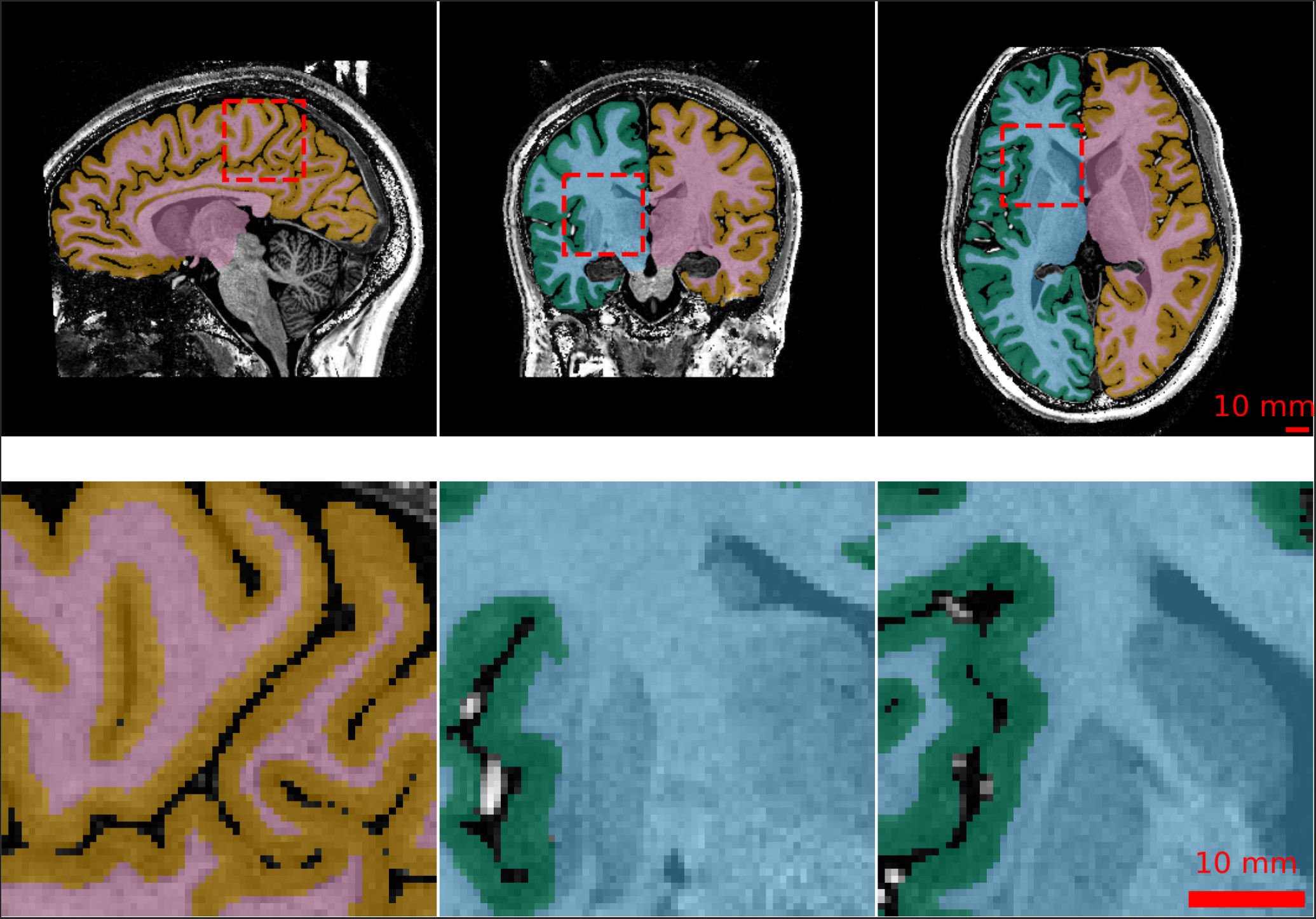
Results of the manually corrected cortical segmentation of Figure 2. Coloured labels are shown in overlay as described in Table 3.

**Figure 4.**
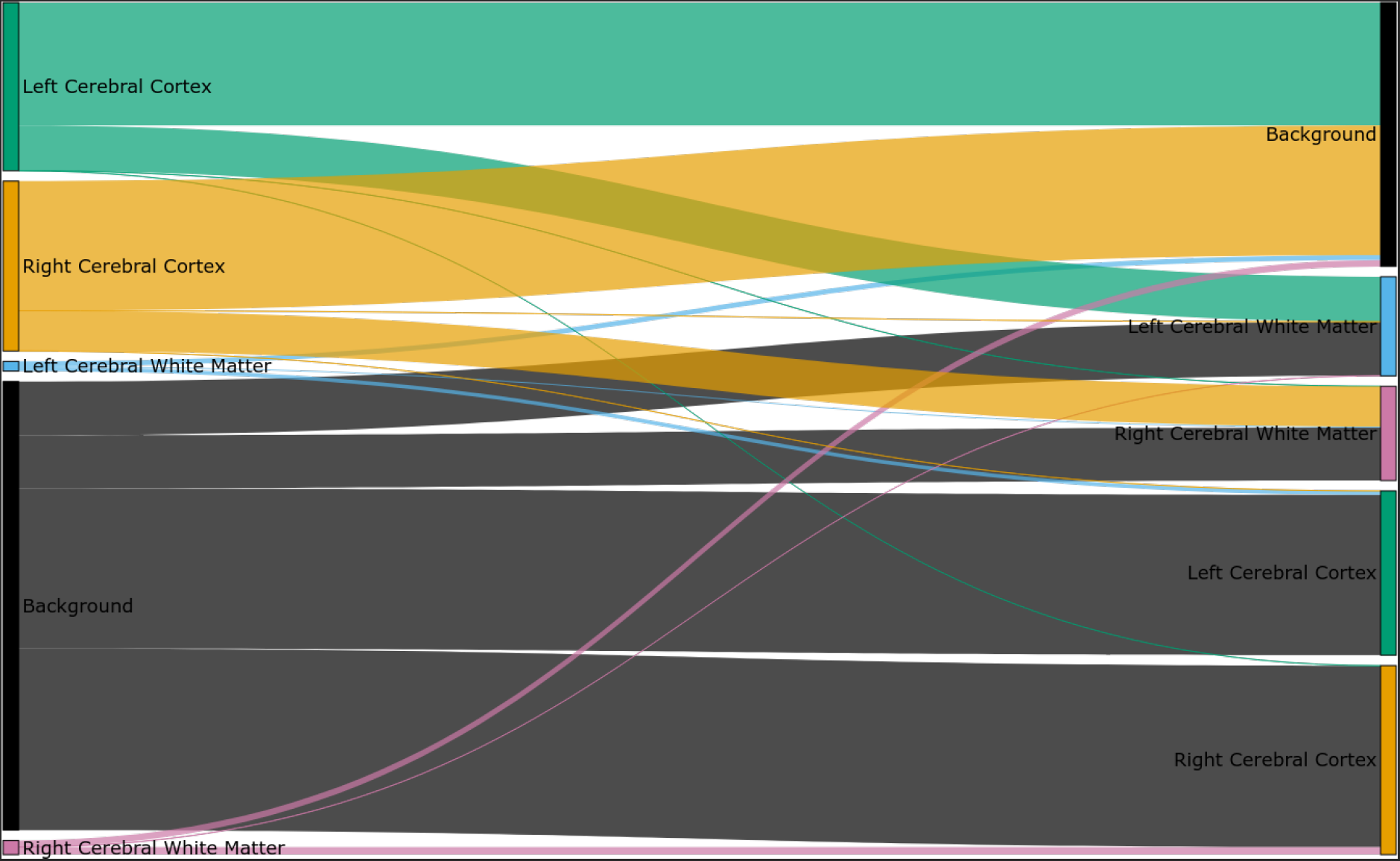
Average change of voxel labels from FreeSurfer (left) to manual segmentation (right) across all subjects. Coloured labels are shown in the sankey diagram as described in Table 3. Cortical labels and background show the largest differences.

### Initial Segmentation Using FreeSurfer

In order to optimize the efficiency of the segmentation process, we used FreeSurfer’s v7.1.1 `recon-all -hires` pipeline^21^ as a preliminary step. However, this approach required adaptations due to the fact that its original design did not account for the high field strengths of 9.4 T and sub-millimeter resolutions *<* 0.8 mm. The initial phase of the `recon-all` pipeline, `-autorecon1`, includes extensive preprocessing steps such as skull stripping, motion correction, intensity normalization, and Talairach transformation computation. Following this phase, the generated brain mask was carefully examined. Should it be necessary, adjustments were made to the pre-flooding height of the watershed algorithm, which is employed to determine a boundary between the brain and the skull. The skull stripping process was repeated until the desired results were achieved. For minor corrections, particularly in the dura mater, the convenient `-gcut` flag was utilized to remove small portions of dura, followed by manual verification of the segmenters.

After these modifications, the manually corrected brain masks were used as input to complete the remaining stages of the `recon-all` pipeline, `-autorecon2 -autorecon3`, using the `-hires` flag. The result was a detailed cortical ribbon mask delineating the segmented volumes of left and right gray and white matter. This ribbon, converted to NIfTI format, served as the basis for the subsequent manual segmentation phase.

### Expert Validation

The manual segmentation procedure commenced with an initial training of the segmenters. Following successful completion, the iterative labeling was performed using ITK-Snap^22^, allowing for voxel-by-voxel corrections in all three anatomical planes. Experienced neuroradiologists and neuroscientists performed regular assessments to verify the accuracy of the tissue delineations. These reviews also served as guidance for the segmenters, providing feedback on the quality of their segmentations and highlighting areas for improvement. Special attention was given to training the segmenters to accurately determine the boundaries between different tissue classes. The iterative process continued until the segmentations were deemed satisfactory by the expert reviewers.

Finally, after the manual corrections were completed and approved in a routine review, the segmentations underwent a final validation phase. This phase involved a thorough inspection in the axial, sagittal, and coronal planes to ensure segmentation consistency, anatomical accuracy, and clear delineation of tissue boundaries. The final validation was conducted by two expert neuroradiologists (B.B., T.L.), who independently reviewed the segmentations. Only those segmentations that successfully passed these stringent validations were considered suitable for inclusion in this publication. This rigorous validation process ensured the high quality and reliability of the segmentations, consistent with the standards required for scientific research in neuroimaging.

## Data Records

### Repository and Dataset Format

The dataset described here is permanently accessible to the public via OpenNeuro^23^ under the accesssion number ds005216 and via the project page https://ultracortex.org. The dataset is released under the Creative Commons license CC0, which allows unrestricted reuse of dataset, including for commercial purposes. The data used in the study were organized using the Brain Imaging Data Structure (BIDS)^24^. Consequently, the associated metadata are stored in files with tab-separated values (TSV) as well as with JavaScript Object Notation (JSON), which include image-specific information about the corresponding demographics and measurement parameters. For each subject we provide the anonymized *T*_1_-weighted image in NIfTI format. Furthermore, we provide the SynthStrip-generated brain mask and skull-stripped image, the FreeSurfer-produced ribbon segmentation that served as the basis for manual segmentation, and the manual segmentation mask as derivative data. The detailed data file directory structure is shown in Figure 5.

**Figure 5.**
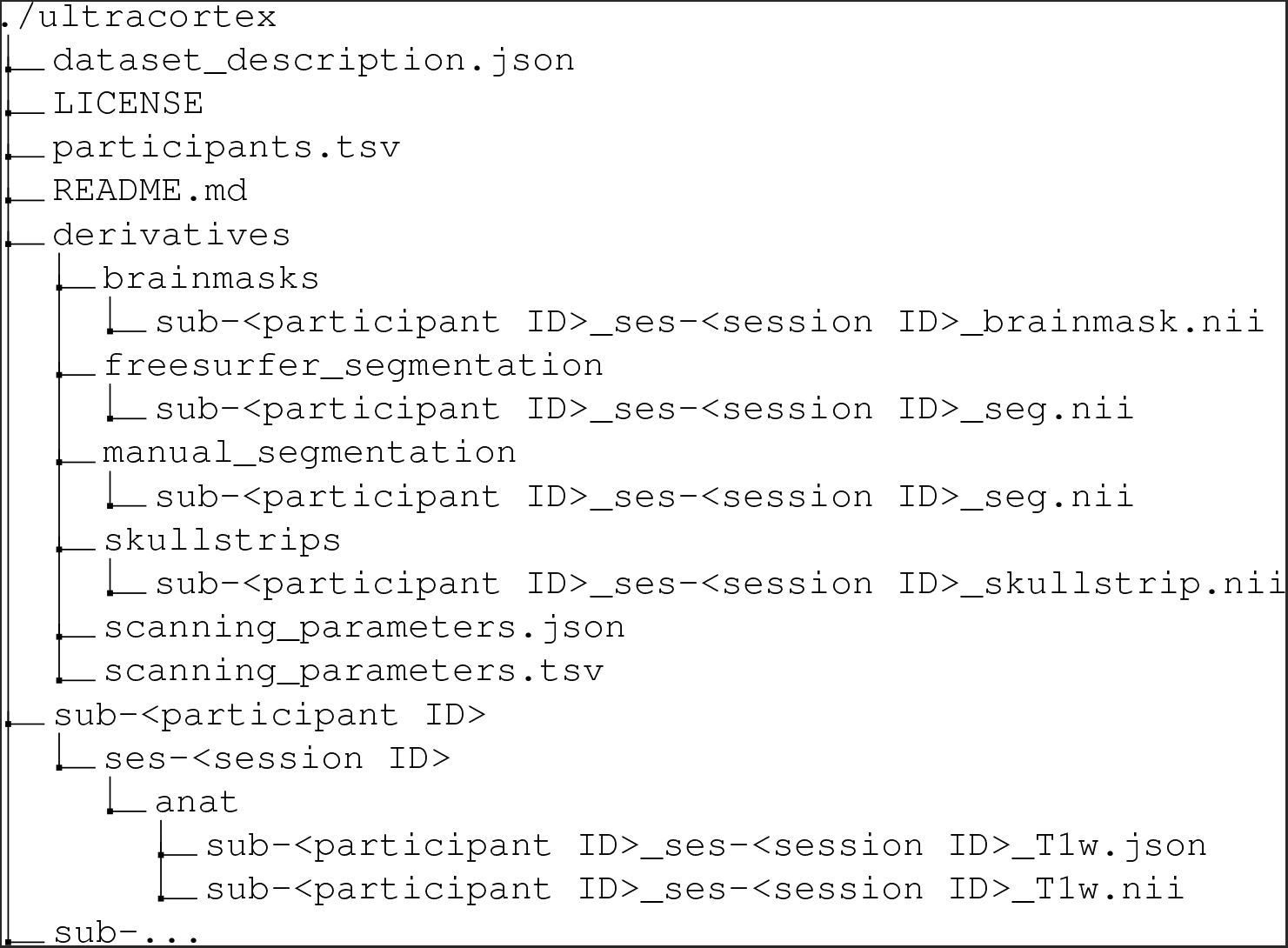
Structure of files and directories.

### Technical Validation

#### Artifacts

The process of evaluating artifacts involved a meticulous, two-tier inspection methodology by MRI experts. Initially, after the acquisition, a qualitative control was conducted, examining the images for any prominent artifacts. 88 images that passed this preliminary inspection were placed into a selection pool. Subsequently, each image within this pool underwent a thorough visual re-inspection, receiving a binary rating based on its quality.

Images containing significant artifacts, such as those resulting from periodic involuntary motion, sudden involuntary movements, conscious body movements, or hardware and sequence-related issues, were excluded from the selection. This process yielded a final dataset of 86 images, with the remaining 2 images excluded due to the presence of significant artifacts.

To quantitatively assess the severity of motion artifacts, we calculated the Entropy Focus Criterion EFC, which leverages the Shannon entropy H of voxel intensities as an indicator of ghosting and blurring due to head motion. The Shannon entropy of an image with *N* voxels is defined by the intensity value *x*_*i*_ of voxel *i* by

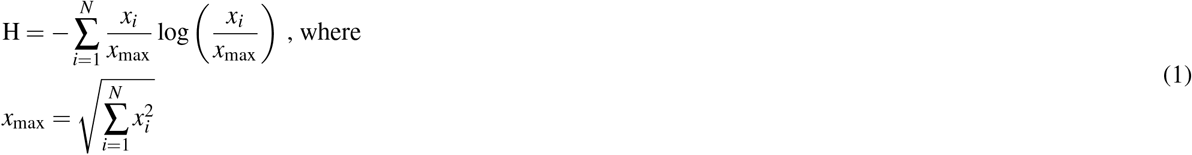

represents the distribution of voxel “brightness”^25^. If all the image energy were concentrated in a single voxel, we would have the greatest possible voxel brightness, *x*_max_. The EFC is defined in accordance with MRIQC^26^ as H normalized by the maximum entropy, which is the case if all voxels have the same value. This implies that 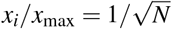, resulting in

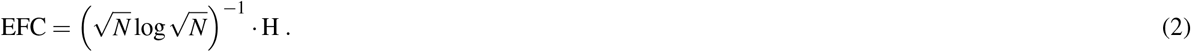

It is preferableto have lower EFC values, with EFC = 0 indicating maximal energy concentration in a single voxel. The mean EFC over all subjects and given sequence configurations is 0.537 *±* 0.057 and a corresponding histogram including kernel density estimate is shown in Figure 6a.

**Figure 6.**
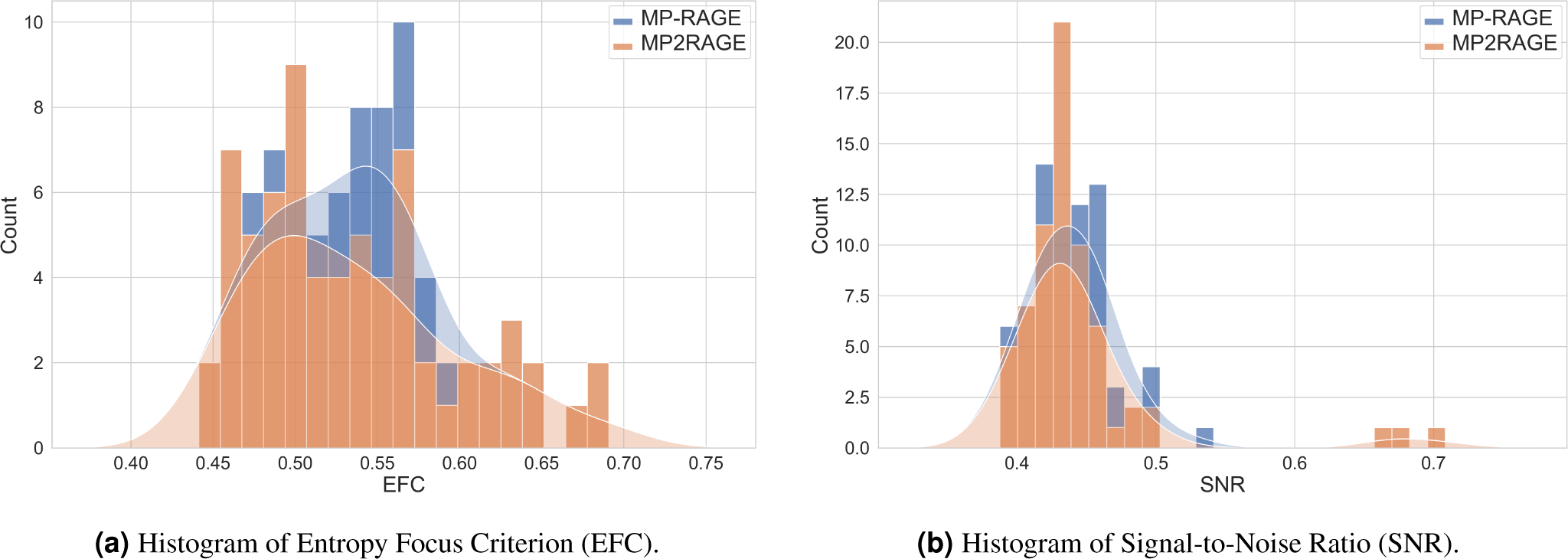
Stacked histograms of image quality metrics with kernel density estimates for MP-RAGE and MP2RAGE sequences.

#### Signal-to-Noise Ratio

To approximate the SNR within the tissue mask, we employed SynthStrip for brain mask extraction. It is important to note that SynthStrip was not initially designed for UHF MRI, particularly not for 9.4 T. Consequently, the resulting skull-stripped image is only an approximate representation 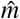 of the actual brain mask *m*. The simplified 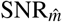 within the approximated brain mask 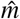 was calculated in accordance with MRIQC by

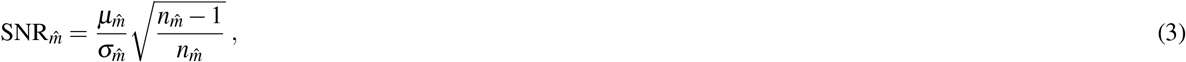

where 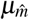 is the mean intensity of voxels within the brain mask, 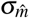 is their standard deviation, and 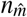 is the total number of voxels in the brain mask. The mean 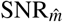 over all subjects is 0.446 *±* 0.053. Figure 6b shows a histogram of 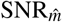 for specific sequence configurations.

#### Segmentations

For the 12 volumes with ground truth segmentations, we conducted additional quantitative evaluations of image quality. Two key metrics were utilized: the Coefficient of Joint Variation CJV and the Contrast to Noise Ratio CNR. The CJV offers insight into the dispersion of tissue intensities relative to their mean intensity and is expressed as follows:

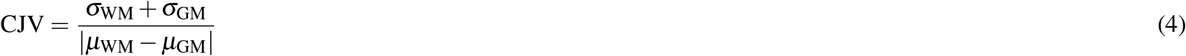

where *σ* _*{* WM,GM*}*_and *µ*_*{*WM,GM*}*_ represent the standard deviation and the mean intensity of all GM and WM voxels, respectively. A lower CJV value indicates a clearer differentiation between tissue classes, with minimal overlap in their intensity distributions.

Furthermore, the CNR metric was employed to assess the difference in intensity between GM and WM regions relative to the background noise. A higher CNR indicates better differentiation between tissue classes. CNR is calculated as:

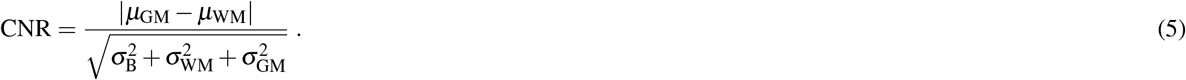

Here, *σ*_B_ denotes the standard deviation of all voxels outside the gray and white matter masks.

The CNR was determinded to be 0.879*±*0.390, while the CJV was 1.683 *±* 1.824. These metrics, CJV and CNR, collectively provide a comprehensive assessment of the segmentation quality in terms of tissue contrast and noise characteristics, thereby ensuring the reliability of the segmentation process in our high-resolution 9.4 T MRI dataset.

### Usage Notes

In this study, we have curated a dataset optimized for automated processing to ensure compatibility and ease of integration into various analytical workflows. The data are provided in widely accepted and well-documented standard formats, including NIfTI and plain text files. This choice of formats facilitates seamless integration into various analytical environments and eliminates reliance on proprietary software. In addition, all data processing for this publication was performed using open-source software on conventional computer workstations. This further enhances the accessibility and reproducibility of our methods.

## Code Availability

All processing pipeline scripts are openly available for download. The code used to generate processed outputs can be accessed via GitHub at https://github.com/MPI-Neuroinformatics/UltraCortex. Further, open-source software packages have been used: dcm2niix multiple versions v1.0.<X> (https://github.com/rordenlab/dcm2niix), FreeSurfer v7.1.1 (https://surfer.nmr.mgh.harvard.edu), ITK-Snap v3.8.0 (http://itksnap.org), Quickshear v1.1.0 (https://github.com/nipy/quickshear), and SynthStrip as provided by FreeSurfer v7.1.1 (https://surfer.nmr.mgh.harvard.edu/docs/synthstrip).

## Acknowledgements

We would like to express our sincere gratitude to all student assistants who played an invaluable role in creating the manual segmentations for this project. Their meticulous efforts, dedication, and attention to detail were critical to the successful completion of this task. We deeply appreciate their hard work and dedication, which greatly enhanced the overall quality and reliability of our research. We gratefully acknowledge funding by the Max Planck Society. This work was supported by the German Research Council (DFG) Grant GZ: GR 833/11-1.

## Author Contributions Statement

Participant recruitment, data acquisition, and qualitative data control: J.B., F.B., E.C., M.E., G.H., R.H., V.K., P.M., and D.R.; Conception, data collection, qualitative data control, and cleaning: J.S.; Design: D.R., L.M., and J.S.; Data processing and anonymization: L.M. and J.S.; Technical validation: L.M.; Supervision of the segmentation process: D.R. and L.M.; Feedback to segmentors and final validation of segmentation: B.B. and T.L.; Writing of the manuscript: L.M. and J.S.; Supervision, participation in discussions and suggestions for manuscript improvement: B.B., G.L., and T.L.; Resources, supervision, and funding acquisition: K.S.; All authors contributed to the article and approved the submitted version.

## References

1. Forstmann, B. U. et al. Multi-modal ultra-high resolution structural 7-Tesla MRI data repository. Sci. Data 1, 140050, DOI: 10.1038/sdata.2014.50 (2014).

2. Alkemade, A. et al. The Amsterdam Ultra-high field adult lifespan database (AHEAD): A freely available multimodal 7 Tesla submillimeter magnetic resonance imaging database. NeuroImage 221, 117200, DOI: 10.1016/j.neuroimage.2020.117200 (2020).

3. Svanera, M., Benini, S., Bontempi, D. & Muckli, L. CEREBRUM-7T: Fast and Fully-volumetric Brain Segmentation of 7 Tesla MR Volumes [Data set], DOI: 10.25493/RF12-09N (2020).

4. Pohmann, R., Speck, O. & Scheffler, K. Signal-to-noise ratio and MR tissue parameters in human brain imaging at 3, 7, and 9.4 tesla using current receive coil arrays. Magn. Reson. Medicine 75, 801–809, DOI: 10.1002/mrm.25677 (2016).

5. Van Essen, D. C. et al. The WU-Minn Human Connectome Project: An overview. NeuroImage 80, 62–79, DOI: 10.1016/j.neuroimage.2013.05.041 (2013).

6. Klein, A. & Tourville, J. 101 Labeled Brain Images and a Consistent Human Cortical Labeling Protocol. Front. Neurosci. 6, DOI: 10.3389/fnins.2012.00171 (2012).

7. Kuijf, H. J. et al. MR Brain Segmentation Challenge 2018 Data, DOI: 10.34894/E0U32Q (2024).

8. Charyasz, E. et al. Functional mapping of sensorimotor activation in the human thalamus at 9.4 Tesla. Front. Neurosci. 17, DOI: 10.3389/fnins.2023.1116002 (2023).

9. Shajan, G. et al. A 16-channel dual-row transmit array in combination with a 31-element receive array for human brain imaging at 9.4 T. Magn. Reson. Medicine 71, 870–879, DOI: 10.1002/mrm.24726 (2014).

10. Mugler III, J. P. & Brookeman, J. R. Rapid three-dimensional T1-weighted MR imaging with the MP-RAGE sequence. J. Magn. Reson. Imaging 1, 561–567, DOI: 10.1002/jmri.1880010509 (1991).

11. Marques, J. P. et al. MP2RAGE, a self bias-field corrected sequence for improved segmentation and T1-mapping at high field. NeuroImage 49, 1271–1281, DOI: 10.1016/j.neuroimage.2009.10.002 (2010).

12. Hagberg, G. et al. Whole brain MP2RAGE-based mapping of the longitudinal relaxation time at 9.4T. NeuroImage 144, 203–216, DOI: 10.1016/j.neuroimage.2016.09.047 (2017).

13. Hurley, A. C. et al. Tailored RF pulse for magnetization inversion at ultrahigh field. Magn. Reson. Medicine 63, 51–58, DOI: 10.1002/mrm.22167 (2010)._eprint: https://onlinelibrary.wiley.com/doi/pdf/10.1002/mrm.22167.

14. Griswold, M. A. et al. Generalized autocalibrating partially parallel acquisitions (GRAPPA). Magn. Reson. Medicine 47, 1202–1210, DOI: 10.1002/mrm.10171 (2002).

15. Breuer, F. A. et al. Controlled aliasing in parallel imaging results in higher acceleration (CAIPIRINHA) for multi-slice imaging. Magn. Reson. Medicine 53, 684–691, DOI: 10.1002/mrm.20401 (2005).

16. O’Brien, K. R. et al. Robust T1-Weighted Structural Brain Imaging and Morphometry at 7T Using MP2RAGE. PLOS ONE 9, e99676, DOI: 10.1371/journal.pone.0099676 (2014).

17. Li, X., Morgan, P. S., Ashburner, J., Smith, J. & Rorden, C. The first step for neuroimaging data analysis: DICOM to NIfTI conversion. J. Neurosci. Methods 264, 47–56, DOI: 10.1016/j.jneumeth.2016.03.001 (2016).

18. Hoopes, A., Mora, J. S., Dalca, A. V., Fischl, B. & Hoffmann, M. SynthStrip: skull-stripping for any brain image. NeuroImage 260, 119474, DOI: 10.1016/j.neuroimage.2022.119474 (2022).

19. Schimke, N. & Hale, J. Quickshear Defacing for Neuroimages. In Proceedings of the 2nd USENIX conference on Health security and privacy, HealthSec’11, 11 (USENIX Association, USA, 2011).

20. Fischl, B. et al. Whole Brain Segmentation: Automated Labeling of Neuroanatomical Structures in the Human Brain. Neuron 33, 341–355, DOI: 10.1016/S0896-6273(02)00569-X (2002).

21. Zaretskaya, N., Fischl, B., Reuter, M., Renvall, V. & Polimeni, J. R. Advantages of cortical surface reconstruction using submillimeter 7 T MEMPRAGE. NeuroImage 165, 11–26, DOI: 10.1016/j.neuroimage.2017.09.060 (2018).

22. Yushkevich, P. A. et al. User-guided 3D active contour segmentation of anatomical structures: Significantly improved efficiency and reliability. NeuroImage 31, 1116–1128, DOI: 10.1016/j.neuroimage.2006.01.015 (2006).

23. Mahler, L. et al. “UltraCortex: Submillimeter ultra-high field 9.4t brain mr image collection and manual cortical segmentations”, DOI: doi:10.18112/openneuro.ds005216.v1.0.1 (2024).

24. Gorgolewski, K. J. et al. The brain imaging data structure, a format for organizing and describing outputs of neuroimaging experiments. Sci. Data 3, 160044, DOI: 10.1038/sdata.2016.44 (2016).

25. Atkinson, D., Hill, D., Stoyle, P., Summers, P. & Keevil, S. Automatic correction of motion artifacts in magnetic resonance images using an entropy focus criterion. IEEE Transactions on Med. Imaging 16, 903–910, DOI: 10.1109/42.650886 (1997).

26. Esteban, O. et al. MRIQC: Advancing the automatic prediction of image quality in MRI from unseen sites. PLOS ONE 12, e0184661, DOI: 10.1371/journal.pone.0184661 (2017).

